# A novel coalescent method insensitive to population structure

**DOI:** 10.1101/2020.11.30.405431

**Authors:** Zeqi Yao, Kehui Liu, Shanjun Deng, Xionglei He

## Abstract

Conventional coalescent inferences of population history make the critical assumption that the population under examination is panmictic. However, most populations are structured. This complicates the prevailing coalescent analyses and sometimes leads to inaccurate estimates. To develop a coalescent method unhampered by population structure, we perform two analyses. First, we demonstrate that the coalescent probability of two randomly sampled alleles from the immediate preceding generation (one generation back) is independent of population structure. Second, motivated by this finding, we propose a new coalescent method: *i*-coalescent analysis. *i*-coalescent analysis computes the instantaneous coalescent rate (iCR) by using a phylogenetic tree of sampled alleles. Using simulated data, we broadly demonstrate the capability of *i*-coalescent analysis to accurately reconstruct population size dynamics of highly structured populations, although we find this method often requires larger sample sizes for structured populations than for panmictic populations. Overall, our results indicate *i*-coalescent analysis to be a useful tool, especially for the inference of population histories with intractable structure such as the developmental history of cell populations in the organs of complex organisms.

## Introduction

Coalescent theory is widely used to infer the demographic history of populations (Drummond et al., 2005; Li and Durbin, 2011; Liu and Fu, 2015; Strimmer and Pybus, 2001). Conventional coalescent inference, e.g. Kingman’s *n*-coalescent, uses the waiting time between coalescent events to estimate effective population size (Kingman, 1982). Such inferences often require the critical assumption that the population under examination is panmictic. Inferences of population history with violated assumption can lead to highly inaccurate estimations (Chikhi et al., 2010; Leblois et al., 2006; Mazet et al., 2016; Wakeley, 1999). For example, while Li and Durbin (2011) suggest that several population size reductions occurred in recent human history, Mazet et al. (2016) show that population structure could explain these signals. Because most populations are not panmictic, it is often necessary to incorporate population structure into coalescent models. This can greatly increase model complexity and requires the use of much larger datasets to make reliable statistical inferences (Grusea et al., 2019; Peter et al., 2010). Furthermore, if structural information is intractable, model construction and data collection become infeasible. These complications necessitate a coalescent method that is insensitive to population structure.

In this paper, we present a new coalescent method to circumvent the influence of population structure, *i*-coalescent analysis. In contrast to other methods which focus on the waiting time between coalescent events (such as Kingman’s *n*-coalescent), *i*-coalescent analysis computes the instantaneous coalescent rate (iCR) of every generation. Mathematical deductions and computational simulations show that the *i*-coalescent analysis can be used to reconstruct the dynamics of population size for highly structured populations.

## Results

We first present two key results: (1) population structure influences coalescent analysis and (2) coalescent probability is independent of population structure for the immediate preceding generation (*t =* 1). Motivated by the latter result, we derive a novel nonparametric coalescent approach, *i*-coalescent analysis, that is insensitive to population structure.

We consider a group of haploid individuals of population size *N* (equivalent to a diploid group of size *N*/2) under a classical *n*-island model. The sub-population of each island evolves as a Wright-Fisher process with no migration allowed between islands. We first consider two simple scenarios (*A* and *B*). In scenario *A, n* = 1. Therefore, the entire population is panmictic with no structure. In scenario *B, n* = 2 and the size of the two sub-populations are *N*_1_ and *N*_2_, respectively, in which *N*_1_ + *N*_2_ = *N*. According to standard coalescent theory in a discrete-generation model, for scenario *A*:

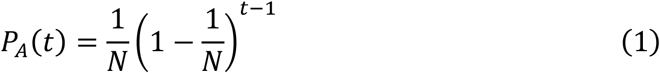

where *P*_*A*_(*t*) is the probability that two alleles randomly sampled from the population coalesce at generation *t*. Note that *t* = 0 depicts the present, and larger *t* indicates further back in the past. Similarly, in scenario *B*:

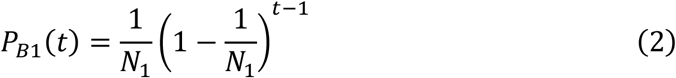

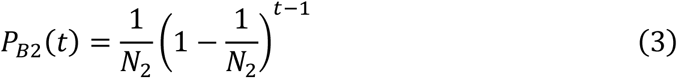

where *P*_*Bi*_(*t*) is the probability that two alleles sampled randomly from the *i*_th_ island coalesce at generation *t*, where *i =* 1, 2. Accordingly, for two alleles sampled randomly from the two islands, the coalescent probability at generation *t* is

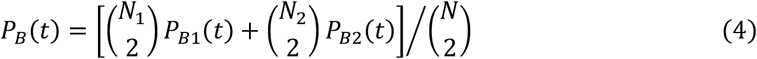

For large *N*, equation (4) can be approximated by

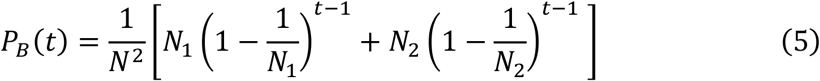

The difference between *P*_*A*_(*t*) and *P*_*B*_(*t*) (equations (3) and (5)) highlights the influence of population structure on coalescent analysis.

Importantly, for *t* = 1, *P*_*A*_ = *P*_*B*_ = 1/*N*. However, the difference between *P*_*A*_(*t*) and *P*_*B*_(*t*) increases with increasing *t* for all combination of *N*_1_ and *N*_2_ (Fig. 1a). Therefore, the coalescent probability in scenario *B* is independent of population structure only at the immediate preceding generation (*t =* 1).

**Fig. 1.**
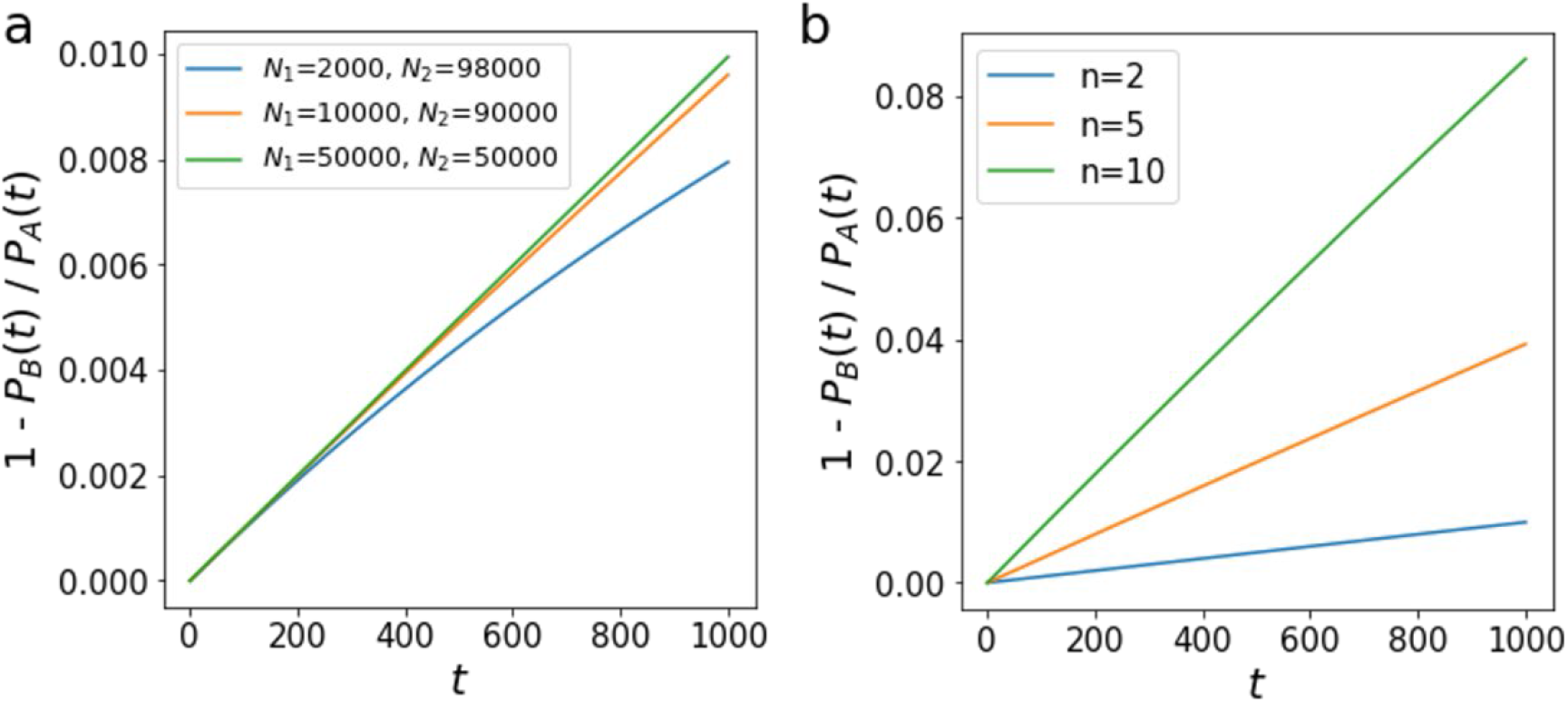
The difference between *P*_*A*_(*t*) and *P*_*B*_(*t*) under a constant-size model. Figures depict a population consisting of *n* constant-size sub-populations. Total population size *N* = 100,000 in all cases. **a**. The curve of 1 - *P*_*A*_(*t*)/*P*_*B*_(*t*) where *n* = 2 with different combinations of *N*_1_ and *N*_2_. **b**. The curve of 1 - *P*_*A*_(*t*)/*P*_*B*_(*t*) where *n* = 2, 5, 10, respectively, with equal sub-populations sizes (*N*_*i*_ *= N/n* for sub-population *Bi*).

We obtain similar results for when n > 2 (a greater number of islands and, consequently, a higher complexity of the population structure). Extending equation (5) to scenarios with n > 2 yields

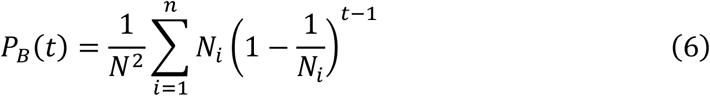

where *N*_*i*_ is the size of each sub-population, *B*_*i*_. For *t* > 1, the difference between *P*_*A*_(*t*) and *P*_*B*_(*t*) increases as *n* increases (Fig. 1b). Thus, the impact of population structure on coalescent analysis increases with the former’s complexity. However, the key observation is that *P*_*A*_(*t*) and *P*_*B*_(*t*) are always identical for *t* = 1, irrespective of the number of sub-populations. This again demonstrates that population structure has no impact on the coalescent probability on the immediate prior generation under the constant-size model.

We extent these results to the exponential-growth model, in which

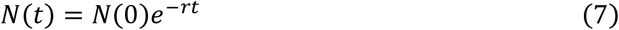

where *N*(*t*) is the size of population at generation *t, N*(0) is the population size at the present, and *r* is the population growth rate. Under this model, *P*_*A*_(*t*) and *P*_*B*_(*t*) can be calculated by the following expressions

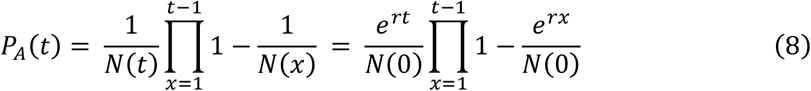

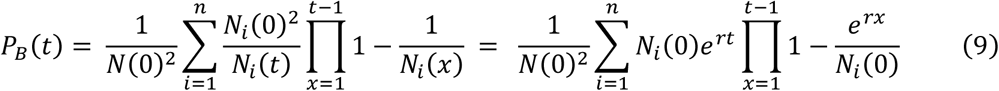

Similar to the constant-size model, the difference between *P*_*A*_(*t*) and *P*_*B*_(*t*) increases with *t* under different initial population sizes in the two-island case (*n =* 2; Fig. 2a) and under different values of *n* (Fig. 2b). Also according to the previous case, the difference between *P*_*A*_(*t*) and *P*_*B*_(*t*) increases with increasing population structure complexity (greater *n*; Fig. 2b). Most importantly, *P*_*A*_(*t*) and *P*_*B*_(*t*) are identical if and only if *t* = 1, assuming each sub-population has the same growth rate. Thus, as in the constant-size model, the coalescent probability at the first prior generation under an exponential-growth model is insensitive to population structure.

**Fig. 2.**
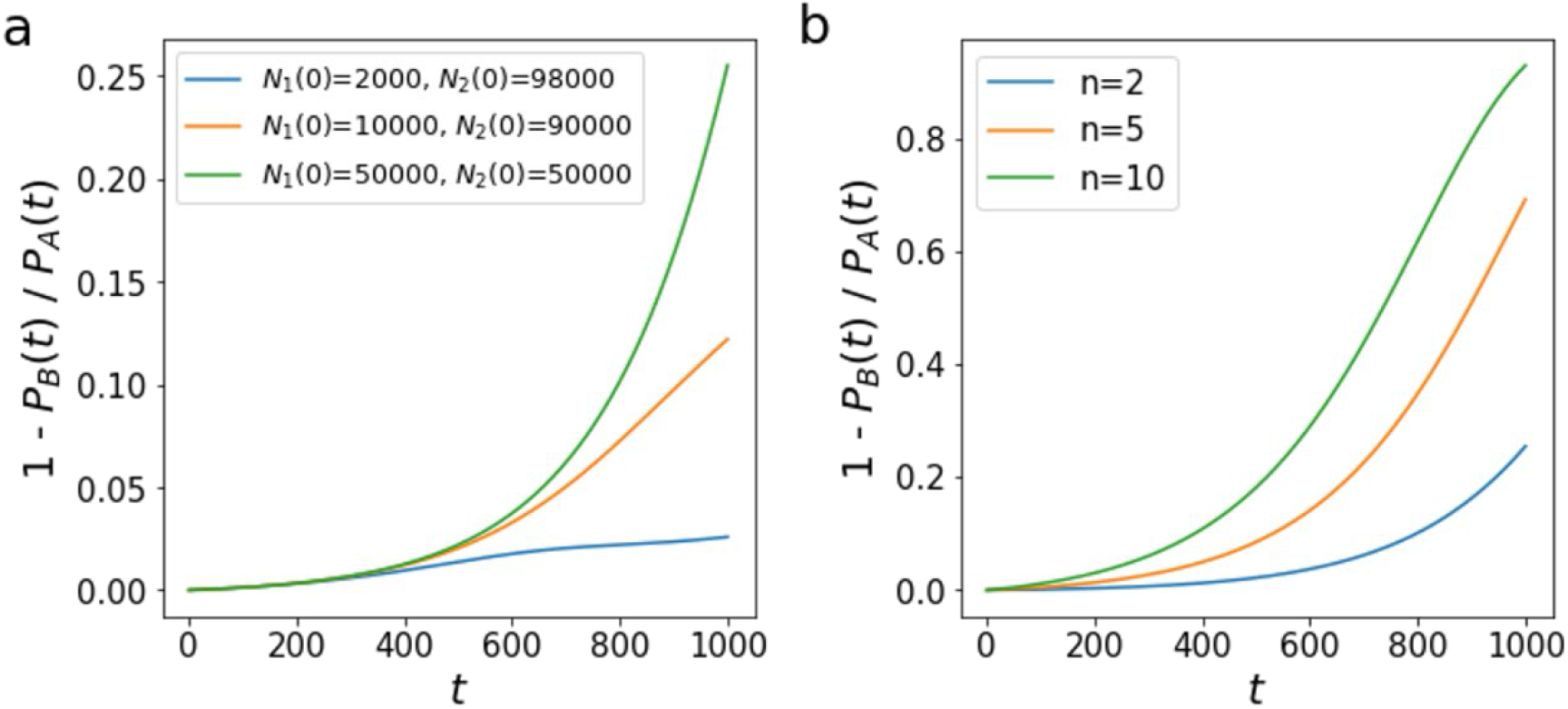
The difference between *P*_*A*_(*t*) and *P*_*B*_(*t*) under an exponential-growth model. Figures depict a population consisting of *n* sub-populations experiencing exponential-growth: *N*(*t*) *= N*(0)*e*^*-rt*^, with *r* = 0.005 for each sub-population and *N*(0) = 100,000 for the whole population. **a**. The curve of 1 - *P*_*A*_(*t*)/*P*_*B*_(*t*) where *n* = 2 with different *N*_1_(0) and *N*_2_(0) combinations. **b**. The curve of 1 - *P*_*A*_(*t*)/*P*_*B*_(*t*) where *n* = 2, 5, 10, respectively. Each sub-population has initial sizes (*N*_*i*_(0) *= N*(0)*/n* for sub-population *Bi*).

The above analysis indicates that an inference method for population history based on the coalescent probability of two randomly sampled alleles at the immediate prior generation, henceforth the instantaneous coalescent rate (iCR) may be insensitive to population structure. Therefore, we develop a novel nonparametric coalescent approach, *i*-coalescent analysis, that estimates the population size of single generation by computing the iCR. With a sufficiently large sample size, the iCR of each generation can be calculated from a phylogenetic tree using the formula

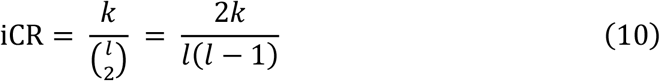

where *k* is the number of coalescent events and *l* is the number of lineages at the examined generation on the phylogenetic tree. Accordingly, the population size *N* can be estimated as

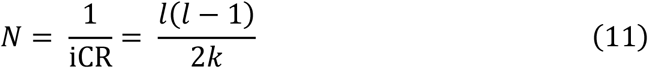

noting that *N* is also instantaneous, corresponding to the specific generation, *t*, examined. See **Methods** for more details.

To validate the efficacy of *i*-coalescent analysis, we carried out simulations. We simulated two populations: (1) a panmictic population, *A*, of haploid individuals and (2) a structured population, *B*, consisting of two sub-populations, *B*_1_ and *B*_2_ (Fig. 3, **Methods**). Populations *A* and *B* were parametrized to be with the same overall size dynamics for 100 generations, in which *B*_1_ and *B*_2_ had distinct growth patterns. After the population growth simulations, 1%, 5%, 10% of the final individuals were sampled, respectively, to reconstruct phylogenetic trees for *i*-coalescent analysis (**Methods**). For population A, *i*-coalescent analysis was able to accurately reconstruct population size dynamics, even with a sampling proportion of 1% (Fig. 3a). For population *B, i*-coalescent analysis yielded accurate results with medium (5%) and large (10%) sample sizes (Fig. 3b). Therefore, *i*-coalescent analysis is capable of providing accurate population size estimates in structured populations, although structured populations require larger sample sizes than their panmictic counterparts.

**Fig. 3.**
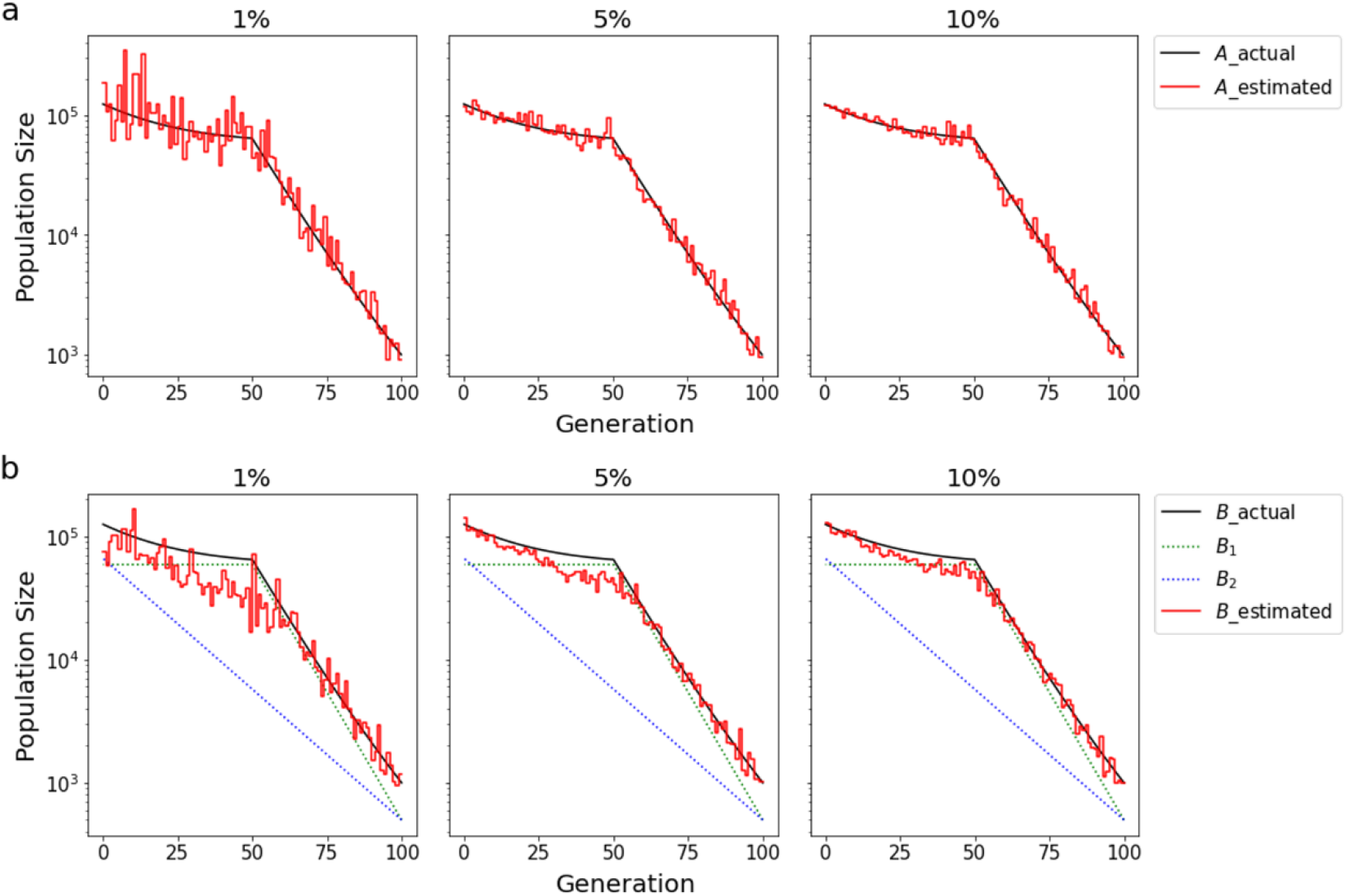
Inference of population size dynamics for a simulated population consisting of two sub-populations. **a**. A simulated panmictic population, *A*. The demographic history is depicted with solid black line. **b**. A simulated structured population, *B*, consisting of two sub-populations *B*_1_ and *B*_2_. The simulated demographic history of each sub-population is distinct from each other, as depicted by the dashed lines. The demographic history of the whole population *B* is the same as that of *A*. The solid red curve in each subplot represents the estimated population size dynamics, which were obtained using *i*-coalescent analysis. Estimated population size dynamics were inferred using samples of 1%, 5%, 10% of the final population, as shown above each subplot.

We also considered a scenario with a more complex structured population, consisting of five sub-populations with distinct demographic histories (Fig. 4, **Methods**). Using *i*-coalescent analysis to infer size dynamics of the simulated populations, we obtained similar results to those shown in Fig. 3. Interestingly, *i*-coalescent analysis performance in the complex scenario (Fig. 4b) was approximately as accurate as in the simpler case (Fig. 3b). Thus, *i*-coalescent analysis can reconstruct population size dynamics for highly structured populations.

**Fig. 4.**
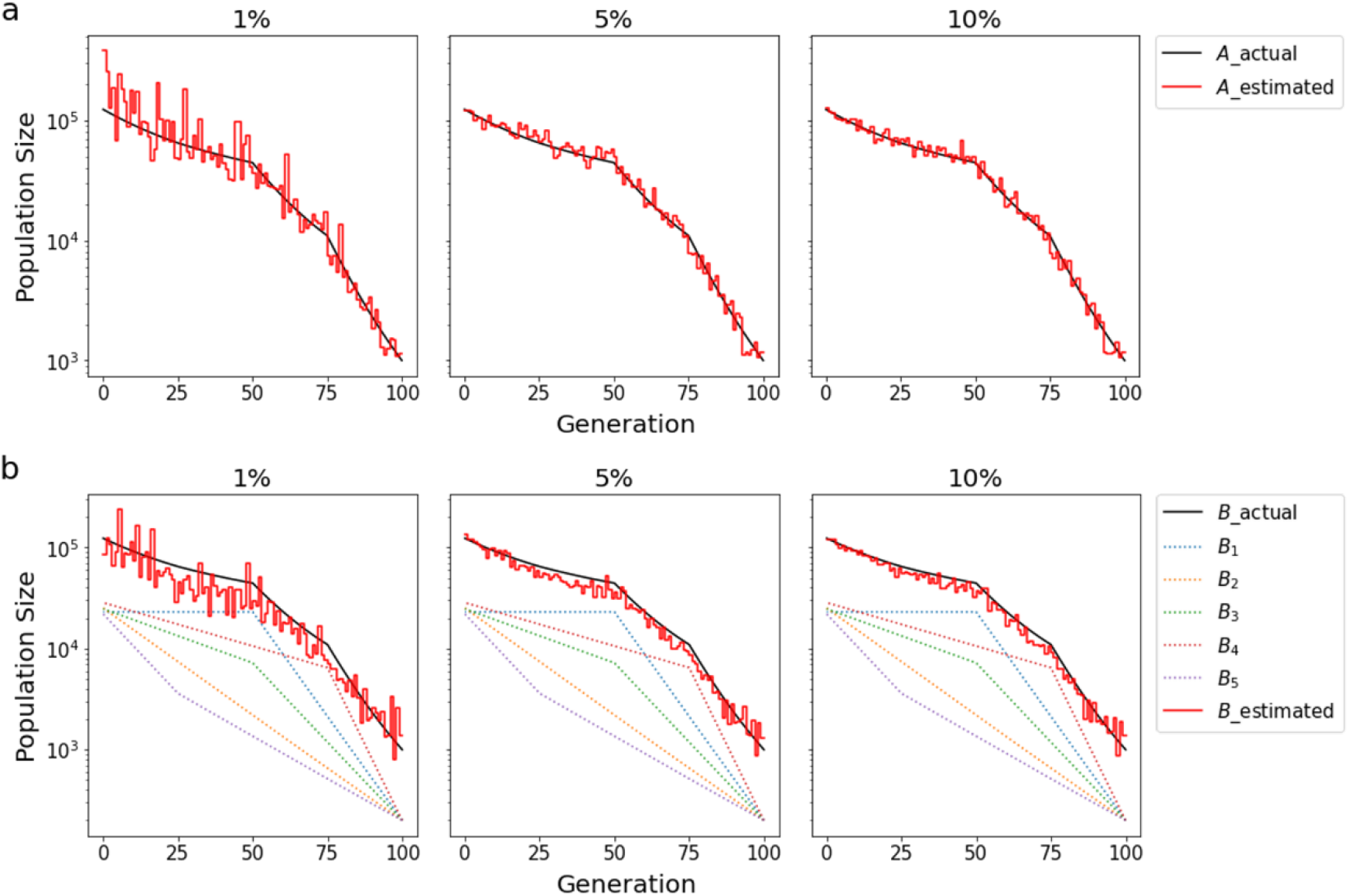
Inference of population size dynamics for a simulated population consisting of five sub-populations. **a, b**. The same simulations as that shown in Fig. 3, except that the simulated structured population (*B*) in **b** consists of five sub-populations, each with distinct demographic histories.

## Discussion

Population structure often hampers efforts to perform coalescent analysis. In this study, we propose a new coalescent method, *i*-coalescent analysis, to infer the dynamics of population size by computing the instantaneous coalescent rate (iCR) of every single generation. Motivated by the result that the coalescent probability of two randomly sampled alleles at the immediate prior generation is insensitive to population structure, *i*-coalescent analysis provides a coalescent method insensitive to population structure. We use simulations to validate this method and show that *i*-coalescent analysis is capable of making reliable inferences of the population size dynamics of highly structurally complex populations.

A necessary prerequisite of *i*-coalescent analysis is that the sample size must be sufficiently large such that the iCR of every generation of a phylogenetic tree can be estimated (see **Methods**). However, the increasing capacity of DNA sequencing has led to an accumulation of data, and nearly one million sequenced human genomes available (Bycroft et al., 2018). Therefore, this sample size requirement does not pose an insurmountable obstacle for the application of this method in making population genetics inferences. Another potential application of *i*-coalescent analysis is the study of cell population history in the organs of multicellular organisms (Hu et al., 2013; Lee-Six et al., 2018). It is well-known that the development of an organism is a process consisting of cell population growth and differentiation. Because the cell populations are highly structured, conventional coalescent methods are incapable of making reliable inferences regarding developmental history. We believe that the *i*-coalescent analysis will be a useful tool for reconstructing organ development when cell phylogeny can be mapped by the use of developing cell barcoding techniques.

## Methods

### Inference of population size dynamics using *i*-coalescent analysis

For each generation *t*, the instantaneous population size is estimated as the inverse of instantaneous coalescent rate (iCR), using the formula

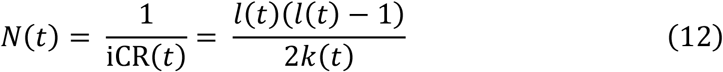

where *N*(*t*) is the instantaneous population size at generation *t*, iCR(*t*) is the instantaneous coalescent rate at generation *t, l*(*t*) is the number of examined lineages at generation *t*, and *k*(*t*) is the number of coalescent events at generation *t*. The complete estimated population size dynamics are obtained from estimating *N*(*t*) for each generation, from the tips (*t* = 0) to the root of the phylogenetic tree.

We note that if the sample size is insufficiently large, there may be no coalescent event at a given generation *t* (*k*(*t*) = 0). In this case, iCR(*t*) is equal to 0 and *N*(*t*) cannot be estimated by equation (12). In these cases, we search for the smallest value of *g* such that *k*(*g*) > 0 where *g* > *t*. We then calculate the coalescent rate for time interval [*t, g*] as the mean of iCR of every generation in the interval. Therefore, the population size in time interval [*t, g*] is estimated by the formula

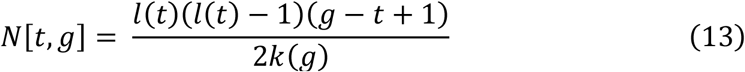

in cases for where *k*(*t*) = 0.

### Simulations under structured and unstructured scenarios

To test the performance of *i*-coalescent analysis, we carried out a set of simulations under structured and unstructured scenarios. We compared an unstructured population to a classical *n*-island model for *n =* 2 and *n =* 5. In each scenario, the initial size of a population of haploid individuals was set 1000. The initial size of each sub-population was set 500 for *n* = 2, and 200 for *n* = 5 where *n* is the number of sub-populations. The growth trajectory of each sub-population was designed to be independent from one another (see the dashed lines in Fig. 3b and Fig. 4b). In unstructured scenarios, a panmictic population was simulated with the same demographic history as the whole population from structured scenarios (see the solid black lines in Fig. 3 and Fig. 4). All simulated populations evolved for 100 generations under a discrete-generation model. At each generation, newborn individuals was randomly assigned a parent from the previous prior generation. Each individual was then assigned a new mutation. To record the genealogical information, no recurrent mutations were allowed. After each simulation, 1%, 5%, 10% of the final individuals were sampled. Each sample was used to reconstruct phylogenies by tracing the genealogy of each individual.

## Acknowledgments

We are grateful to the members of the He lab for comments on the manuscript.

## Author Contributions

H.X., D.S., L.K. and Y.Z. conceived the study; D.S., L.K., Y.Z. and H.X. analyzed data; H.X. supervised the study and wrote the manuscript with inputs from co-authors.

## Author Information

The authors declare no competing interests.

## Notes

### Competing Interest Statement

The authors have declared no competing interest.

